# Adult human *ex vivo* brain slices for dissecting glial biology and multicellular communication

**DOI:** 10.64898/2026.01.12.698855

**Authors:** Miriam Adam, Inbar Shapira, Hanan Schoffman, Gavin Piester, Zhaorong Li, Francisco J. Quintana, Iddo Paldor, Tal Shahar, Naomi Habib

**Affiliations:** Edmond & Lily Safra Center for Brain Sciences, The Hebrew University of Jerusalem, Israel; The Laboratory of Molecular Neuro-Oncology, Shaare Zedek Medical Center, Faculty of Medicine, The Hebrew University of Jerusalem, Jerusalem, Israel; Ann Romney Center for Neurologic Diseases, Brigham and Women’s Hospital, Harvard Medical School, Boston, MA, USA; Department of Pathology and Laboratory Medicine, University of Rochester Medical Center, Rochester, NY 14642, USA; The Gene Lay Institute of Immunology and Inflammation, Brigham & Women’s Hospital, Massachusetts General Hospital and Harvard Medical School, Boston, MA, USA; The Broad Institute of MIT and Harvard, Cambridge, MA, USA; The Department of Neurosurgery, Shaare Zedek Medical Center, Faculty of Medicine, The Hebrew University of Jerusalem, Jerusalem, Israel; The Department of Neurosurgery, Tel Aviv Sourasky Medical Center, Affiliated with Faculty of Medicine, Tel Aviv University, Tel Aviv, Israel

**Author notes:** Equal contribution (Miriam Adam, Inbar Shapira, Tal Shahar, Naomi Habib).

## Abstract

Glial cells are critical modulators of brain function in health, aging, and disease, emerging as promising therapeutic targets. However, exploring their roles and therapeutic potential is limited by the lack of experimental systems that faithfully capture the repertoire of mature human glial cells while permitting controlled perturbations. Here, we establish a robust *ex vivo* platform for cell-type specific interrogation of glial responses and multicellular crosstalk, based on adult human organotypic brain slice cultures, obtained from neurosurgical resections. We show that slice cultures preserve tissue architecture, maintain all major cell types and mature cellular identities over weeks in culture. These cultures elicit robust, stimulus-specific transcriptional programs to diverse stressors and inflammatory stimuli, demonstrating sensitivity to distinguish closely related signals and reproducibility despite biological and technical variation. Moreover, we resolved coordinated glial cell type-specific responses to TNFα, a key mediator of neuroinflammation, uncovering distinct and physiologically relevant functional roles validated in postmortem human brains. Network analyses discovered balanced pro– and anti-inflammatory loops among microglia and astrocyte cells, which notably also involved oligodendrocyte precursor cells (OPCs), confirming their suggested role in regulation of the tissue level inflammatory response in human brains. We experimentally validated that glial activation in slice cultures is orchestrated not only by direct stimulation but also through intercellular signaling across cell types, mirroring natural multicellular dynamics in brain tissue. Together, organotypic brain slice cultures emerge as a sensitive and robust platform for dissecting adult human glial biology, paving the way for deeper mechanistic insights and advanced drug-screening applications.

## Introduction

Deciphering mechanisms underlying brain functions in health and disease and enabling therapeutic discovery, requires experimental systems that faithfully reproduce the cellular and structural complexity of the adult human brain while allowing controlled perturbations. Glial cells are increasingly recognized as key modulators of brain physiology^1,2^ and as primary drivers of various neurodegenerative diseases, neuroinflammation, brain aging and injury^3–5^. Owing to their high plasticity, glial populations were shown to undergo coordinated shifts in cellular states that can either promote neuroprotection or trigger neurotoxicity, thereby shaping disease outcomes^6–11^ – and positioning them as promising therapeutic targets. Unlocking their biological roles and therapeutic potential requires human models that preserve the native milieu of mature glial cells, their physiological functions, and their interactions within adult brain tissue architecture.

Modeling the adult human brain remains a major challenge. Murine models are widely used but are limited by species-specific genetic and functional differences, particularly evident in disease modeling and glial biology^12–14^, reducing their translational relevance and faithful recapitulation of human neuropathologies^15,16^. A significant advancement has been the derivation of brain cells from human embryonic stem cells (ESCs) and induced pluripotent stem cells (iPSCs), enabling detailed perturbations in 2D and 3D culture models. However, harnessing these systems for studies of the adult and aging brains is still limited: differentiation protocols struggle to generate fully mature cells with physiological functions^17–19^, and direct reprogramming approaches, while shown to generate mature-like neurons^20^, are not yet available for glia and retain epigenetic memory of the donor cell. Moreover, robust derivation protocols for several glial types remain incomplete, such as oligodendrocyte precursor cells (OPCs) that are particularly challenging to model *in vitro.* Brain organoids provide improved spatial context and extracellular architecture compared to monolayer cultures, yet they still resemble embryonic-like states with incomplete representation of glial and vascular cells^21^. While efforts to improve these culturing systems continue^22,23^, these limitations underscore the unmet need for a complementary model that preserves adult human brain cells within their native brain architecture.

Organotypic brain slice cultures are a powerful *ex vivo* platform that preserves native cytoarchitecture of brain tissue. Established initially in postnatal mice^24^, these cultures have been extended to adult human brains derived from neurosurgical resections of both healthy brain tissue^25–28^ and brain tumors^29–31^. This system bridges the gap between human *in vitro* cell models and adult human brains, as it can facilitate manipulations and long-term investigations of brain physiology, disease mechanism and therapeutic applications. However, prior work in adult human organotypic slice cultures has focused primarily on neuronal viability, physiology and connectivity^26–28^, while their utility for glial biology has not been systematically explored. Key questions therefore remain: Can this platform be leveraged to probe human glial functions? Are mature glial milieu, physiological responses, and intercellular interactions preserved in adult brain slice cultures?

Here, we establish adult human organotypic brain slice cultures as a platform for systematic investigation of human glial biology and multicellular crosstalk. We show that slice cultures preserve tissue architecture and mature cellular diversity over extended culture duration, with glia populations exhibiting robust, reproducible, and stimulus-specific physiological responses, mirroring postmortem human brain signatures. By exposing the slice cultures to a key neuroinflammatory cytokine TNFα, we resolve glial cell type-specific neuroinflammatory responses, providing the first demonstration of coordinated multicellular signaling in live human brain tissue. Network analyses revealed TNFα-driven signaling from neighboring cells that collectively shape glial activation; this signaling is validated in perturbed iPSC-derived astrocytes and consistent with crosstalk observed in intact postmortem brain tissues, establishing the physiological relevance of the platform. Notably, we found balanced pro– and anti-inflammatory loops among microglia and astrocytes, and identified OPCs as emerging contributors to tissue-level inflammatory orchestration in human brains. Together, our findings position adult human organotypic brain slice cultures as a unique and powerful platform for dissecting adult human glial functions and multicellular communication, as well as for evaluating their therapeutic potential.

## Results

### Ex-vivo human organotypic brain slice cultures robustly preserve glial niches

To establish the ex vivo organotypic adult human brain slice cultures as an adult human brain 3D model for functional interrogation of multicellular glial responses, we used tumor free resected specimens from cortical regions routinely obtained during common neurosurgical procedures (**Fig. 1a**, **Methods**). Evaluation by immunohistochemistry confirmed cell survival in culture and preservation of overall tissue architecture, with neurons spread throughout the slices, in line with previous reports (see 3D neuronal staining by MAP2 antibody in **Supplementary movie**), and further demonstrated that microglia and astrocytes remained preserved and co-localized with neurons across the brain slices (**Fig. 1b**).

**Figure 1.**
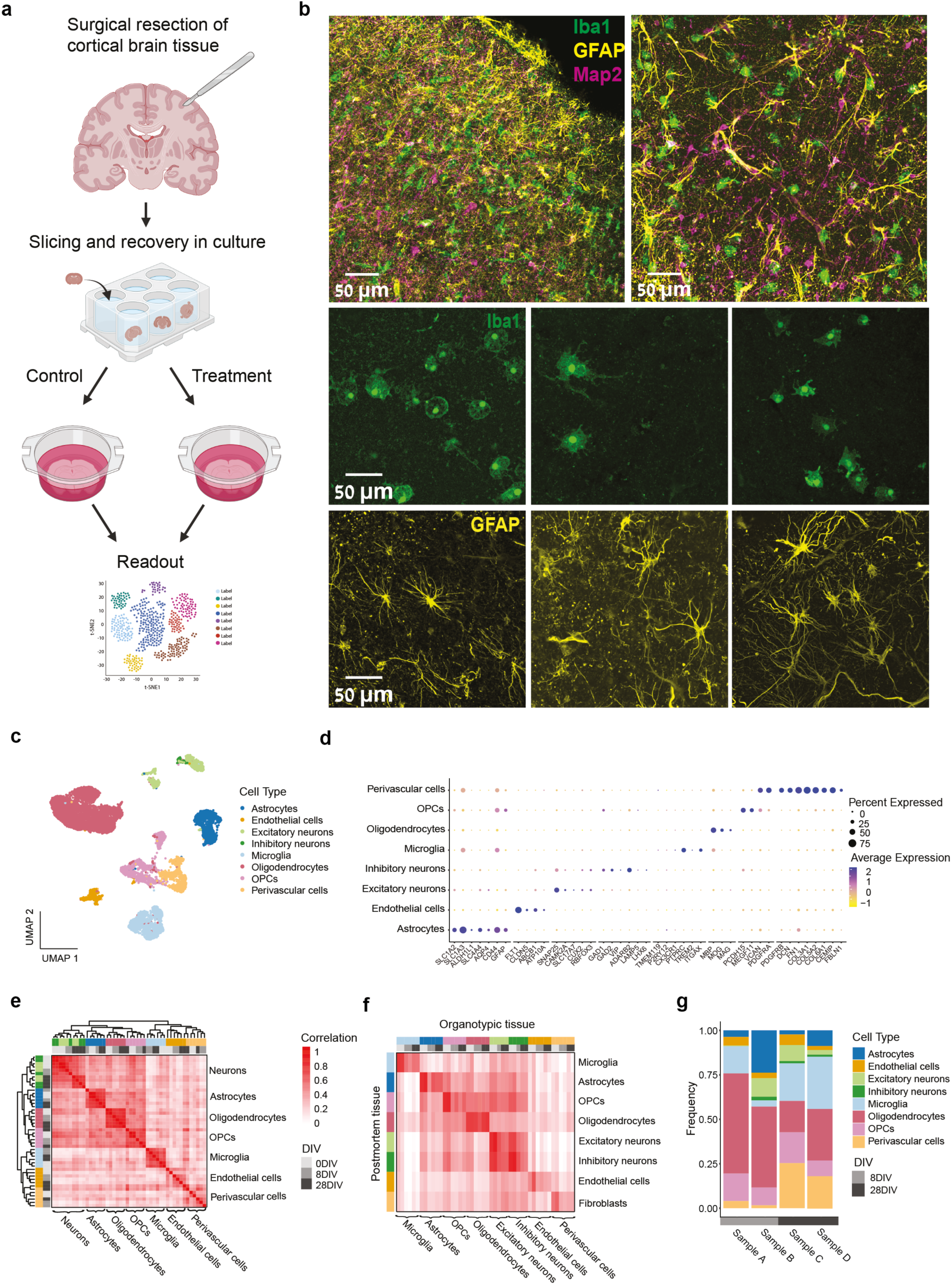
Cellular characterization of organotypic brain cultures. **a**, Overview of the experimental methodology. **b**, Immunofluorescent staining of cultured brain slices with markers for neurons (MAP2, magenta), astrocyte (GFAP, green), and microglia (Iba1, yellow). The composite images are 20x magnification and the individual channels are 40x. **c-d**, Cell type composition in adult brain slice cultures. UMAP embedding of snRNA-seq profiles of ∼20,000 nuclei (n=4, 8 and 28 DIV), colored by cell type (c); Expression levels of selected cell type marker genes across cell types (d). **e-f**, Conserved cell type identities in organotypic slice cultures. Pairwise correlations between cell-type profiles (pseudobulk snRNA expression), comparing cell types across cultured (8/28 DIV) and acute (non-cultured, 0 DIV) slices across donors and culture duration (e); comparing cell types across aging postmortem human brain with cultured and acute brain slices (f). Colorbar: Cell type identity, Days In Vitro (DIV). **g,** Cell type compositions across samples. The relative abundance of cell types per sample.

We characterized the cellular composition of the cultured slices and their ability to preserve the full spectrum of endogenous brain cell populations over extended culturing duration. We applied single-nuclei RNA sequencing (snRNA-seq) to profile cellular compositions in slices from four individuals cultured for 8 or 28 days in vitro (DIV) (**Supplementary Table 1**). Clustering analysis of 11,924 nuclei, resolved all major brain cell types, including excitatory and inhibitory neurons, microglia, astrocytes, oligodendrocytes, OPCs, endothelial cells, and other vascular niche cells (**Fig. 1c**). Cells clustered by cell type, rather than donor or culture duration, indicating that slice cultures preserve cellular identity despite inter-individual variability, effects of tissue handling and time in culture (**Fig. 1c-d, Extended Data Fig. 1a-d**).

To validate that cell identity in culture reflects their state in mature human brain, we compared transcriptional profiles of cultured cells at both culture durations (8 and 28 DIV) with acute resected tissue (matching samples and tissue handling procedures) and a published postmortem human brain snRNA-seq dataset^10^. Cells of the same type showed strong transcriptional correlations across cultured slices, acute tissue, and postmortem brains, with clustering driven by cell type rather than condition. Importantly, cell identity remained stable over time in culture, with similarly high correlations observed at both 8 and 28 DIV (**Fig. 1e-f**). These findings confirm that organotypic slice cultures maintain stable, mature cellular identities comparable to those of the intact human brain.

Assessment of cellular composition (**Fig. 1g**) showed that all brain cell types were preserved across samples and culture duration, though their relative proportions varied. In line with previous reports, neuronal abundance showed substantial variability and declined with longer culture durations, likely reflecting the increased vulnerability of adult human neurons to resection and *ex vivo* culturing^32^. In contrast, glial populations remained stable within sample over time, while their relative proportions varied between individuals, consistent with expected inter-individual differences and technical factors (**Fig. 1e** and **Extended Data Fig.1e**). Together, these findings demonstrate that organotypic slice cultures robustly maintain glial niches over extended culture durations, highlighting their value as a practical and broadly applicable model for investigating glial biology.

### Differential responses to external stimuli in organotypic brain slice cultures

To assess the relevance of the organotypic slice cultures for investigation of physiological responses of adult human brain cells, we exposed the cultured slices to a panel of external stimuli relevant to aging brains. We applied two pro-inflammatory signals: TNF⍺ (cytokine^33^) and LPS (bacterial endotoxin^34^) as well as two stressors: oxidative stress (H_2_O_2_) and a DNA-damaging agent bleomycin (BLM^35^). Each stimulus was applied to samples (n=3-6 per stimulus, at 8 or 21 DIV, from at least 2 individuals) and the responses were profiled by RNA-sequencing 24 hours post stimuli (6 hours for H_2_O_2_) (**Extended Data Fig.2a-c)**. We found that slice cultures showed distinct responses to each stimulus, as revealed by Linear Discriminant Analysis (LDA, controlling for individual-specific differences, **Methods**). Projection of samples into the LDA space clearly separated transcriptional profiles by the stimulus, while samples from different individuals exposed to the same stimulus clustered together, confirming that slice cultures consistently capture coherent, stimulus-specific transcriptional responses (**Fig. 2a**, and **Extended Data Fig.2d**).

**Figure 2.**
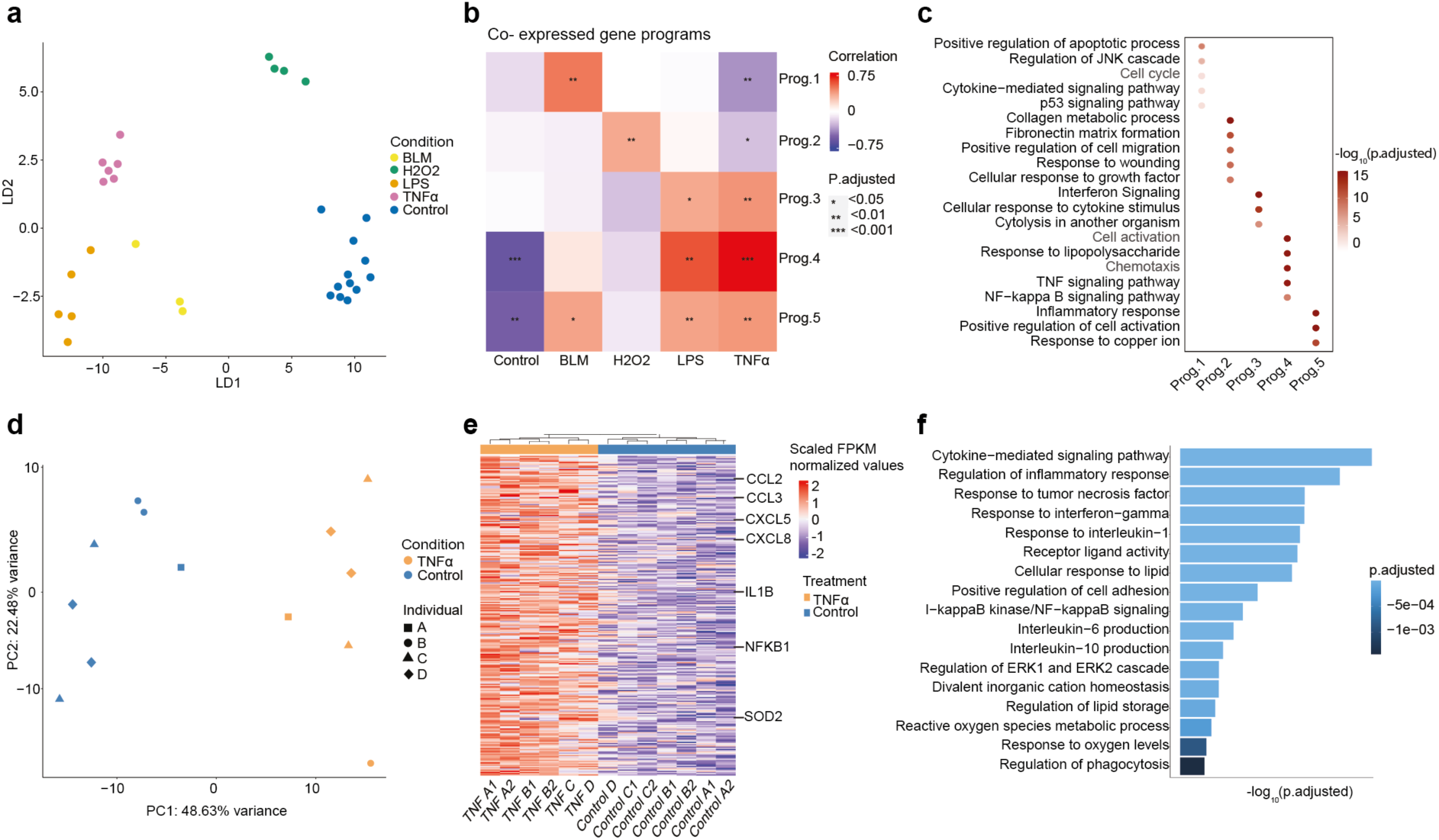
Differential response of adult human organotypic brain slice cultures to stimuli. **a**, Distinct transcriptional responses to stimuli in slice cultures. Embedding of bulk RNA-seq profiles in LDA space, shows differential transcriptional response of slice cultures following TNF⍺, LPS, H₂O₂, BLM (DNA damage)-treatment. **b**, Transcriptional programs linked to stimuli. Selected gene co-expression programs (WGCNA, **Methods**) linked to the different treatments. Corrected p-value (FDR) for correlations between programs and treatments (color scale), FDR corrected p-value: *: *p* < 0.05, **: *p* < 0.01, ***: *p* < 0.001. **c**. Enriched pathways per gene co-expression program. adjusted hypergeometric p-value (FDR)<0.05 (−log_10_P.adjusted). **d**, Robust response to TNF⍺ across slice cultures. Embedding of bulk RNA profiles of TNF⍺ treated and control slice cultures in PCA space. **e**, TNF⍺ induced genes (scaled log_2_ FPKM-normalized counts, log_2_ (fold change) > 1.5, adjusted p-value (FDR) <0.05). **f**. Enriched pathways of significantly TNF⍺ upregulated genes (FDR adjusted p-value (FDR) <0.05).

To further compare responses across the four stimuli, we applied Weighted Gene Co-expression Network Analysis (WGCNA^36^), which identifies sets of co-expressed genes (programs) across samples, without prior knowledge of treatment groups. We identified five gene programs significantly associated with specific treatments: a DNA damage program (associated with BLM, Prog.1), an oxidative stress program (H₂O₂, Prog.2), two inflammatory programs (TNF⍺ and LPS, Prog.3-4), and a broader stress-related program shared across H₂O₂, TNF⍺ and LPS (Prog.5, **Fig. 2b-c**, **Supplementary Table 2 and Extended Data Fig.2f-k**). While TNF⍺ and LPS both engaged immune-related programs, TNF⍺ induced a stronger and more distinct activation (**Extended Data Fig. 2i-j**), highlighting differences in the inflammatory responses. Pathway enrichment analysis annotated each of the programs uncovering stimulus-specific functions that align with previous literature; for example, the DNA damage program (Prog.1) was enriched for apoptosis genes^37,38^, whereas the oxidative stress program (Prog.2) was enriched for wound healing genes^39^. Together, these results demonstrate that organotypic slice cultures capture physiologically relevant stimulus-specific responses, overcome technical and donor variability, and sensitively discriminate between overlapping inflammatory programs and convergent stressor effects.

### TNFα induces robust, coordinated, and cell type-specific responses in brain slice cultures

To dissect the robustness and cell type-coordination of human brain slice cultures in response to perturbations, we investigated the response to TNF⍺, a central mediator of CNS pro-inflammatory signaling and neurodegeneration^40,41^. The well-established transcriptional response to TNF⍺ can be used for validation of our system. Bulk RNA-seq profiles of slices treated with TNF⍺ (4 individuals, n=2 at 8 DIV and n=2 at 28 DIV) confirmed highly reproducible responses across individuals, cortical sub-regions, and culturing duration (**Fig. 2d-e**). Principal component analysis (PCA) showed that TNF⍺ exposure emerged as the dominant driver of transcriptomic profiles, exceeding both biological and technical effects (PC1 = variance explained 48.6%, **Fig. 2d**). Differential expression analysis identified 308 upregulated and 104 downregulated genes robustly modulated across samples and culturing times (FDR adjusted p-value <0.05, log_2_ fold change >1.5; **Fig. 2e, Supplementary Table 3 and Extended Data Fig. 2e**). The TNF⍺-induced genes included the expected canonical inflammatory mediators and hallmarks of TNF⍺ signaling, such as NFKB1 (master regulator^42^), IL1B (pro-inflammatory cytokine^43^), and multiple chemokines (CCL2, CCL3, CXCL5 and CXCL8). Pathway enrichment confirmed activation of inflammatory pathways across cell types (e.g. *cytokine mediated signaling*, *response to TNF*), as well as microglia-associated processes such as lipid storage and phagocytosis (FDR<0.05; **Fig. 2e-f**). The top downregulated genes include microglial AD risk genes (**Extended Data Fig. 2e**): TREM2, a immunomodulatory receptor that was previously shown to be suppressed under pro-inflammatory conditions^36^, and MS4A6A that has a role in neuroprotection and whose downregulation promotes inflammation^44^. Together with downregulation of additional immune response modulators (CD300A^45^ and CD180^38^) these changes indicate the pro-inflammatory shift upon TNF⍺ treatment. Overall, these results underscore the robustness of this system for perturbations and validate the physiological relevance of the response despite potential culture-related effects.

To resolve TNF⍺ responses across cell types, we performed snRNA-seq on human brain slices treated with TNF⍺ or vehicle control (n = 3 individuals; 8 and 28 DIV). The neuronal response to TNF⍺ exposure resulted in reduced viability (**Extended Data Fig. 3d)**, as was shown in previous works^44,45^. The most pronounced transcriptional changes, however, were observed in glial populations (**Fig. 3a**), with NFKB1 consistently upregulated across TNF⍺-treated nuclei, reflecting coordinated activation (**Fig. 3b**). However, the magnitude and nature of responses differed between cell type, with microglia, astrocytes, and OPCs exhibiting the strongest responsiveness. Differential expression analysis (FDR adjusted p-value <0.05, **Methods**) defined cell type-specific signatures of TNF⍺-activation (**Fig. 3d-f**), which were robust across samples and culture duration (**Extended Data Fig. 3a-c).**

**Figure 3.**
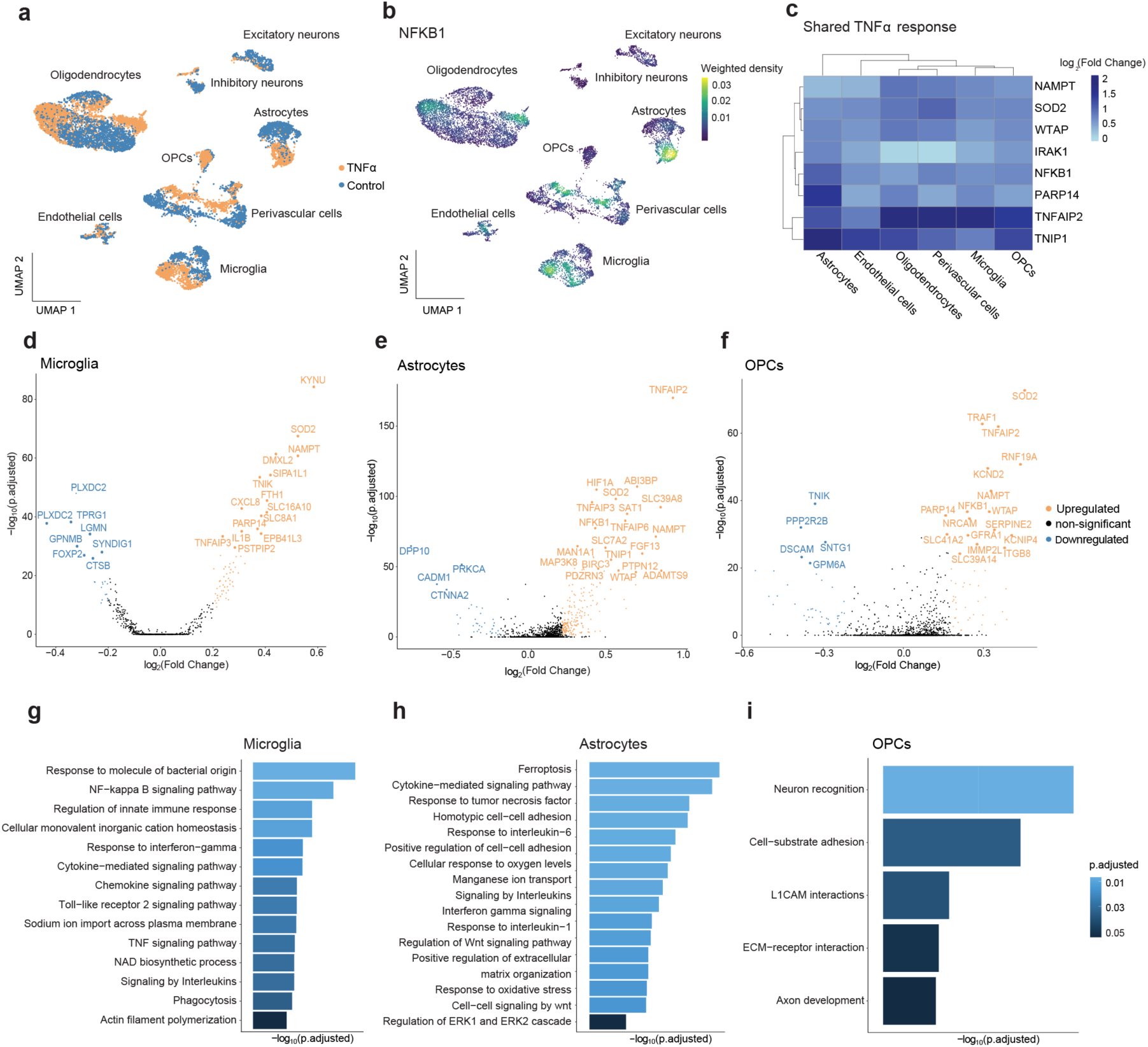
Cell type specific response to TNF⍺. **a**, UMAP embedding of single nuclei transcriptional profiles of TNF⍺ treated and control slice cultures. **b**, NFKB1 expression levels across cell types**c**, Core TNF⍺-induced genes across cell types (log_2_ FC > 0.25, adjusted p-value (FDR) < 0.05). **d-f**, differentially expressed genes in microglia (**d**), astrocytes (**e**) and OPCs (**f**) (log_2_ FC > 0.25, adjusted p-value (FDR) < 0.05). **g-i**, Enriched pathways of TNF⍺ significantly induced genes in microglia (**g**), astrocytes (**h**) and OPCs (i) adjusted p-value (FDR)< 0.05).

Microglia, the brain’s resident immune cells, upregulated NF-κB signaling and pro-inflammatory cytokines (IL1B and CXCL8) known to amplify the initial TNF⍺ signaling^43,46^, while simultaneously inducing negative regulators that limit the inflammatory response (*TNFAIP3*^47^*, PARP14*^48^), indicating an intricate balance between activation and feedback inhibition. They also upregulated adhesion and extracellular matrix interactions (ECM)-related genes (CD44, ITGB8) known to affect microglial inflammatory responses^49,50^, as well as oxidative stress mediators (KYNU^51^, SOD2^52^) and phagocytosis regulators (LYN, SLC11A1 and HCK)^53^ (**Fig. 3d, g**). Together, these define a TNF⍺-activated inflammatory microglial phenotype aligned with the established role of TNF⍺ in neuroinflammation^54^.

Astrocytes also exerted a pro-inflammatory response to TNFα, activating NF-κB signaling (NFKB1, IL33^55^) while simultaneously inducing feedback inhibitors (TNFAIP2^56^, NFKBIA^57^). They also upregulated a broad multifaceted response, including adhesion and ECM molecules (ICAM1, CD38, CD44, FN1), IL6-JAK/STAT signaling (JAK1, JAK2), and autophagy-related genes (MAP1LC3A, ATG7) involved in clearance of pathogens and damaged cells under inflammatory conditions^58^. Elevated HIF1A and PDK3 suggests a metabolic shift under inflammatory stress^59^. Collectively, these changes are consistent with a reactive astrocyte phenotype with both inflammatory signaling and metabolic adaptation^40–42^ (**Fig. 3e, h**).

Beyond microglia and astrocytes, OPCs also exhibited strong inflammatory activation following TNFα, consistent with growing evidence for their role in neuroinflammation in murine models^60,61^, including NF-κB activation (NFKB1), oxidative stress genes (SOD2), and inflammatory-regulators (TNFAIP2, NAMPT). Notably, cell cycle regulator CDK6A^62^ and lineage commitment factor ASCL1^63^ were upregulated, implicating TNFα in potential modulation of OPC differentiation into mature oligodendrocytes (**Fig. 3f, i**).

Oligodendrocytes exhibited more variable responses to TNFα, possibly reflecting sensitivity to varying neuronal survival in cultures and to regional heterogeneity (**Extended Data Fig. 3d**-**e**). Nevertheless, they upregulated known TNFα-induced genes (SOD2, NAMPT, WTAP), similar to other glial cells. The oligodendrocyte response varied with the culture duration, showing upregulation at the earlier timepoint (8DIV) in response to TNFα of myelin-modulating genes (PLLP^64^ DAAM2^65^ and ERBB4^66^), suggesting structural adaptation to TNFα-induced stress.

Moreover, endothelial and peri-vascular cells showed upregulation of inflammatory and stress signatures, shared with glial cells, but not with neurons. These shared genes include a pro-inflammatory core (NFKB1, CD44, NAMPT, and SOD2), potentially contributing to tissue-wide inflammation, which are balanced by the shared expression of anti-inflammatory genes TNIP1 and PARP14 that can dampen the inflammatory response. (**Fig. 3c**).

Collectively, these results demonstrate that slice cultures capture coordinated TNFα-induced programs across all glial cell types, with distinct yet complementary glial responses forming the core of the tissue-wide inflammatory cascade (**Supplementary Table 4 and Extended Data Fig. 4**).

### TNFα-induced responses in brain slices capture physiologically relevant transcriptional programs

We next validated the physiological relevance of the recorded glial responses by comparing their slice-derived transcriptional signatures with external datasets. Microglia and astrocyte responses in slice cultures significantly overlapped with reported up-regulated signatures of genes in TNFα-treated murine microglia^67^ and human iPSC-derived astrocytes^68^, confirming consistency with established models (linear mixed-effects models corrected for individuals, microglia: adjusted p-value (FDR) =5.9e-190; astrocytes: adjusted p-value (FDR) =1.9e-211; **Fig. 4a-b and Extended data Fig. 5a-b**).

**Figure. 4.**
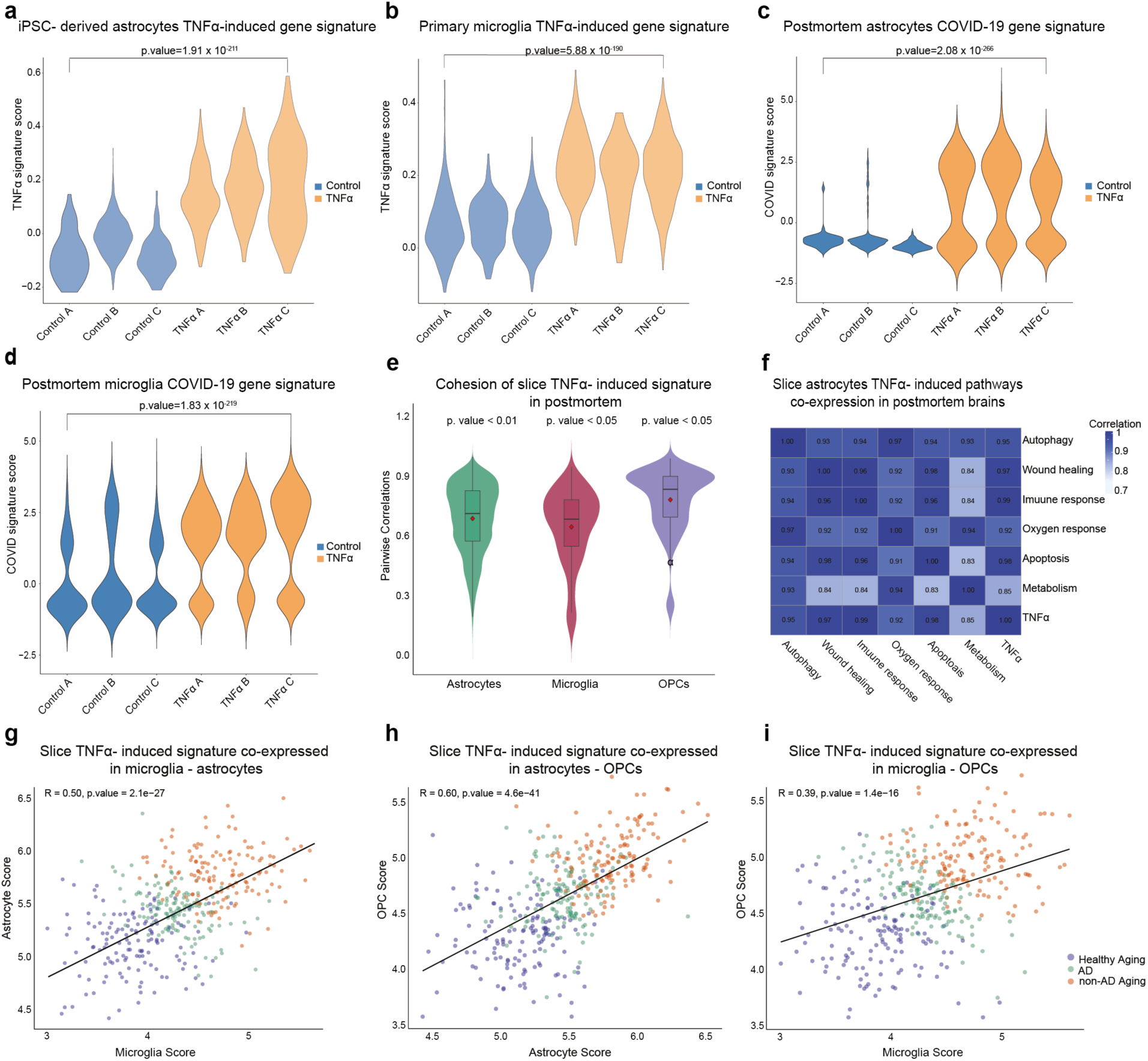
Slice culture TNFα responses recapitulate physiological transcriptional programs. **a-b**, Slice culture TNFα response validation against *in-vitro* transcriptional programs. Distribution of human iPSC astrocytes TNF⍺ signature score across astrocytes of TNF⍺ treated and control cultured slices **(a)**. Distribution of mouse microglia TNF⍺ signature score across microglia of TNF⍺ treated and control cultured slices **(b)** (linear mixed-effects models corrected for individuals, log_2_ FC > 1.5, adjusted p-value (FDR) < 0.05). **c-d**, Slice culture TNFα response validation against COVID19 induced transcriptional changes in astrocytes and microglia from human postmortem brains. Distribution of COVID19 signature scores across astrocytes **(c)** and microglia **(d)** of TNF⍺ treated and control cultured slices (linear mixed-effects models corrected for individuals, log_2_ FC > 0.25, adjusted p-value (FDR)< 0.05). **(e-i)**, Slice culture TNFα response validation against large human postmortem brain cohort (n=437). Co-expression of TNF⍺ signature genes within cell types in post mortem brains. Pairwise gene-gene Spearman correlations were calculated from gene expression of the slice cultures derived TNF⍺ signature genes. Statistical significance was determined by comparing observed correlations with those from 10,000 randomly sampled gene sets of equal size, assessing the probability that TNF⍺-associated gene correlations exceed those expected by chance **(e)**. Pathways upregulated in astrocytes of TNF⍺ treated slice cultures, showing co-expression in post mortem brain astrocytes (Spearman correlation of gene sets annotated to specific functions) **(f)**. **g-i**, Pairs of glial cells from human post mortem brains, show coordinated expression of slice culture TNF⍺ induced genes. Spearman correlations in microglia-astrocytes (**g**), astrocytes-OPCs (**h**), and microglia-OPCs (**i**). Each point represents an individual postmortem sample, color-coded by clinical condition (healthy aging, Alzheimer’s disease, or non-AD aging).

To extend validation to the adult human brain and test if we captured responses of mature glial cells within their native multicellular context, we compared the slice-derived signatures to snRNA-seq datasets from postmortem adult human brain tissue. We first utilized transcriptomic profiles of astrocytes and microglia from postmortem brains infected by the COVID-19 virus, previously shown to capture an acute inflammatory response^69^. Up-regulated gene signatures in COVID-19 brains, in comparison to matched controls, were also significantly up-regulated in TNFα-treated microglia and astrocytes in slice cultures (**Methods**, microglia: adjusted p-value (FDR)= 1.83E-219; astrocytes: adjusted p-value (FDR)=2.08E-266; **Fig. 4c-d and Extended data Fig. 5c-d**).

While informative, the COVID-19 dataset is small and reflects severe acute pathology. Thus, we leveraged a larger cohort of snRNA-seq profiles of 437 aged cortical samples spanning normal aging and Alzheimer’s disease (AD)^10^, both known to associate with chronic inflammation and particularly to TNFα signaling. Indeed, this atlas includes distinct subpopulations of aging astrocytes and microglia exhibiting inflammatory signatures, including TNFα-response genes, providing a reference to validate at scale both the physiological relevance and multicellular coordination of TNFα responses observed in slice cultures^10^. We thus validated that slice-derived TNFα-signatures (up-regulated genes) formed cohesive programs in postmortem aging astrocytes, microglia, and OPCs, *i.e.* the signature genes were significantly co-expressed in each glial cell type (empirical permutation test; adjusted p-values (FDR): astrocytes = 0.006, microglia = 0.022, OPCs = 0.03; **Fig. 4e and Extended data Fig. 5e-g**). Furthermore, the multifaceted astrocytic response to TNFα observed in slice cultures, was confirmed in postmortem brains. We validated the significant co-expression of the different TNFα slice-induced astrocyte pathways in astrocytes from postmortem brains, including, immune activation, autophagy, and metabolic reprogramming, confirming the complex nature of astrocytic TNFα-induced functions (**Fig. 4f**).

Next, we confirmed that these slice-derived TNFα-signatures were coordinated across glial cell types in postmortem tissues: scoring TNFα slice-derived signatures within each cell-type per individual, revealed significant pairwise correlations of the TNFα scores between glial cell types in human brains (astrocytes-OPCs R = 0.6, astrocytes-microglia R = 0.5, microglia-OPCs R = 0.39) (**Fig. 4g-i**). Moreover, stratifying postmortem samples by clinical condition showed that TNFα slice-derived signatures were minimal in healthy aging controls, enriched in typical Alzheimer’s disease (AD), and most pronounced in amyloid-β-pathology associated aging individuals (non-AD alternative aging, characterized by amyloid-β pathology but limited/no neurofibrillary tangle burden^10^). This links the slice-derived TNFα signatures to coordinated glial responses in aging human brains, associated with AD but also with alternative aging processes.

Taken together, these concordant findings validate that our organotypic slice cultures not only capture reproducible, cell-type specific TNFα programs across glial populations but also faithfully recapitulate physiologically relevant and coordinated glial activation patterns observed in adult human brain tissue following viral infection, aging and AD.

### Cell-cell communication orchestrates the glial TNFα-response in slice-cultures

Given the coordinated response across cell types, we next asked how cell-cell communication synchronizes microglia, astrocytes and OPCs activation in slice cultures. We applied NicheNet^70^, which integrates differential gene expression data with known ligand-receptor interactions and their downstream signaling pathways, to predict signaling pathways that are uniquely induced by TNFα compared to controls. TNFα stimulation significantly increased cell-cell communication, predominantly driven by microglia-astrocyte interactions, with substantial crosstalk with OPCs (**Fig. 5a and Supplementary Table 5**).

**Figure 5.**
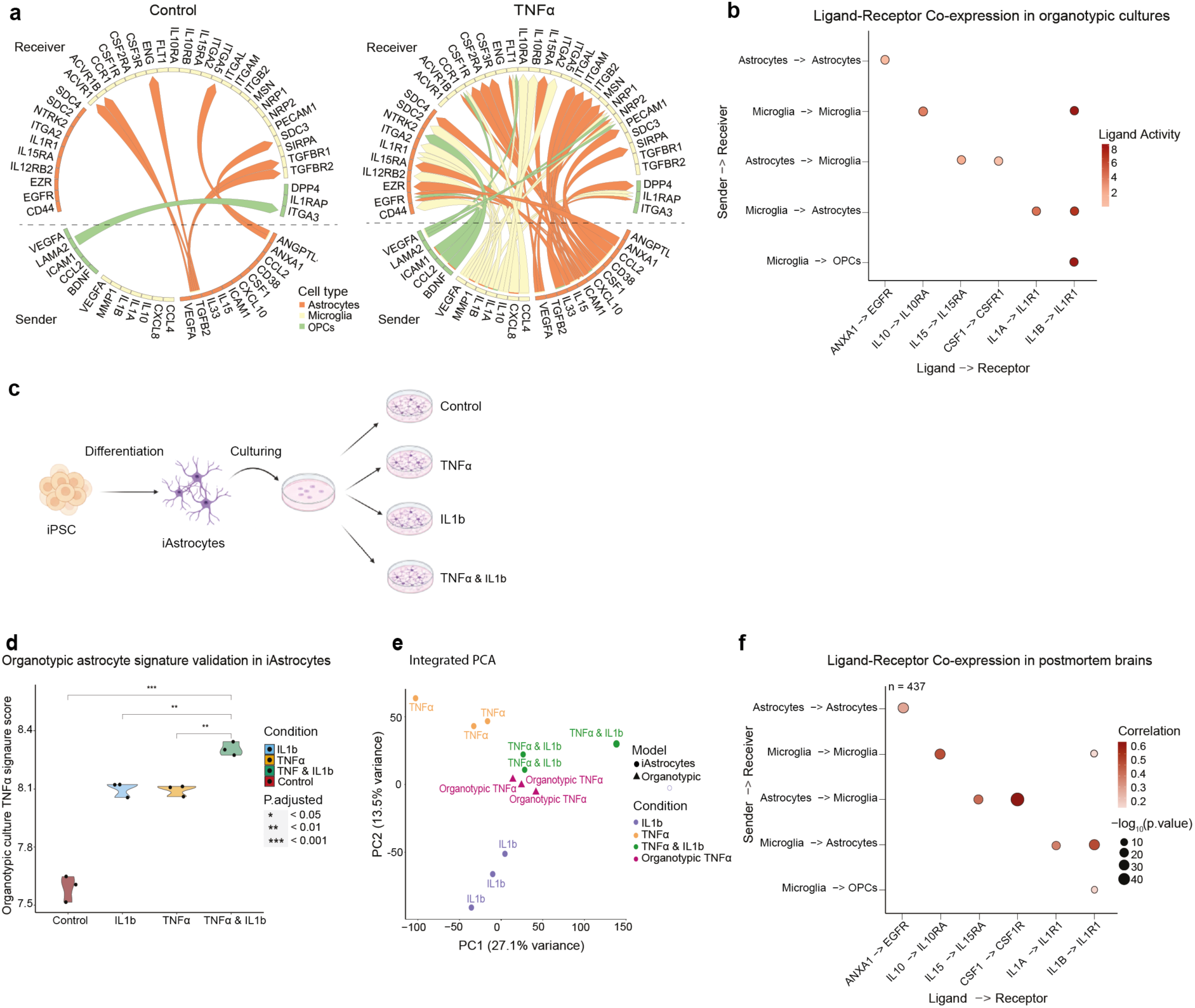
Intricate cellular crosstalk shapes TNF⍺ response. **a**, Predicted differential ligand-receptor interactions in control (left) and TNF⍺-treated (right) slice cultures, as inferred by MultiNicheNet analysis. The width of each arrow reflects the strength of predicted signaling activity. **b,** Selected ligand-receptor pairs across sender-receiver cell type combinations in slice cultures. **c**, Illustration of iPSC treatment experiment **d**, Distribution of TNF⍺ signature scores across iPSC-derived astrocyte exposed to various treatments. Scores were calculated from differentially upregulated genes in TNF⍺ treated slice culture astrocytes (pairwise t-tests with FDR correction). **e,** Integrated PCA plot of transcriptional profiles of iPSC-derived astrocytes and organotypic astrocytes. **f**, Selected ligand-receptor co-expression in human postmortem brains.

Examining the predicted cell-cell communication networks, the microglial transition to a pro-inflammatory state is accompanied by cytokine expression (IL1A, IL1B) that signal to astrocytes, OPCs, and microglia, inducing downstream activation through the NF-κB pathway. Astrocytes, in turn, amplify microglial and OPC inflammatory responses by secreting cytokines and chemokines. For example, astrocytes produce the known pro-inflammatory cytokine IL15 that signals to microglia via IL15RA, leading to NF-κB^71^ pathway activation. Astrocytes also support microglial survival and proliferation through CSF1-CSF1R signaling^72^. In addition, astrocytes and OPCs secrete CCL2 chemokine, which facilitate migration and recruitment of microglia^73^. Finally, ICAM-integrin mediated signaling spans microglia, astrocytes, and OPCs, likely contributing to adhesion migration and communication across the TNFα-treated tissues^74^ (**Fig. 5a-b**).

In parallel to the inter-cell type (paracrine) pro-inflammatory signaling, we identified several intra-cell type (autocrine) anti-inflammatory interactions following TNFα treatment. These included autocrine IL10-IL10R signaling in microglia, a known mechanism to restrict excessive inflammation and restore microglia homeostasis^75^. In turn, astrocytes exhibited autocrine ANXA1-EGFR signaling following TNFα stimulation, a potent anti-inflammatory pathway^76^ that might suppress astrocyte-mediated inflammation (**Fig 5a-b**).

The predicted cell-cell communication network indicates that TNFα-induced changes in slice cultures extend beyond direct stimulation, incorporating secondary inputs from neighboring cells. To test this, we focused on microglia-derived IL-1β signaling to astrocytes, a pathway strongly predicted by our network. Human iPSC-derived astrocytes (iAstrocytes) were treated by TNFα, IL-1β, or both, and profiled by RNA-seq (**Fig. 5c**, n=3 per condition, with PBS controls). Quantifying TNFα slice culture signatures in each iAstrocyte treatment condition, we found the highest expression of the TNFα slice culture signature in the combined TNFα+IL-1β condition, reflecting synergistic activation of astrocytes by both cytokines in the slices (adjusted p-values (FDR)=0.0004 (TNFα+IL-1β), 0.022(TNFα), 0.03(IL-1β); **Fig. 5d**). Integration of the RNA profiles of the slice culture astrocytes and iAstrocytes showed that TNFα-treated slice culture astrocytes clustered closely with TNFα+IL-1β-treated iAstrocytes, while the transcriptional profile of iAstrocytes treated by TNFα-only or IL-1β-only diverged from that of the slice culture astrocytes and from each other (**Methods**, **Fig. 5e**). Together, these results demonstrate that the astrocytes in slice cultures capture a composite of direct stimulus (TNFα) and secondary inputs from neighboring cells (*e.g.* microglial IL-1β), thereby reflecting a physiologically integrated tissue-level response.

Finally, to validate the predicted cell-cell communication network in human brains we examined the coordinated expression of key ligand-receptor pairs across cell types in the 437 postmortem aging brains^10^. Core ligand-receptor pairs identified in our TNFα-treated slice cultures were significantly co-expressed in postmortem brains across the relevant cell types, confirming that the orchestration of glial responses observed in brain slice cultures mirrors TNFα-driven glial communication in human post mortem brain tissue (**Fig. 5b, f**).

## Discussion

In this study, we address a major gap in brain research by establishing adult human organotypic brain slice cultures as a robust *ex vivo* platform for interrogating glial biology and multicellular dynamics in adult human brains. This system provides a unique advantage for studying adult-specific phenotypes and therapeutic discovery for age-associated disorders. By demonstrating reproducible stimuli– and cell-type-specific glial responses that mirror physiological signatures in human brains, we position organotypic slices as a reliable model for cellular and molecular studies of human glial cells.

To our knowledge, this is the only experimental system that enables functional analysis of mature human glia within their native cellular environment and three-dimensional tissue architecture. Other *in vitro* methods, including human ESC/iPSC-derived cultures and organoids, that enable controlled perturbation studies at scale, are challenged by incomplete maturation, lack of representation of certain glial subtypes (*e.g.*, OPCs), ongoing presence of progenitor cells, and potential residual epigenetic memory in derived cells that may affect cellular identity. In contrast, we showed that organotypic slice cultures preserve mature native cellular identity, with the full spectrum of brain cell types, including diverse glial cells with naturally abundant OPCs as well as vascular niche cells, cultured in intact 3D tissue architecture that supports multicellular signaling. Thus, providing a highly needed complementary research platform for translational studies.

Functionally we showed that slice cultures sensitively capture stimulus-specific glial programs, including responses to various stressors, and discriminate between closely related signals such as TNFα and LPS. The measured transcriptional responses are highly reproducible across samples and individuals, despite biological and technical variation. Following TNFα stimulation, we resolved coordinated yet cell type specific programs across astrocytes, microglia, and OPCs, highlighting their complementary roles in neuroinflammation. These signatures aligned with postmortem human brain tissue (COVID response, aging, and AD), underscoring the physiological relevance of the measured responses in slice cultures. Network analyses further revealed that glial activation in slice cultures is shaped not only by direct stimulation but also by secondary inputs from neighboring cells, validated in perturbed iPSC-derived astrocytes and consistent with crosstalk observed in intact postmortem brain tissues.

Our work also revealed the delicate balance of pro– and anti-inflammatory signaling within multicellular brain tissues: microglia propagated TNFα-driven cytokine signaling to astrocytes and OPCs, while astrocytes amplified and diversified responses by integrating TNFα with microglia-derived signaling (e.g. IL1β), in line with previous studies^62,63^. OPCs, often overlooked in neuroinflammation, emerged as active participants and regulators within the glial network, consistent with recent reports implicating them in regulation of brain homeostasis in mice^77^ and in balancing natural aging and AD^78^. Moreover, cell-cell communication analyses highlighted how inter-cell-type paracrine pro-inflammatory signaling (*e.g.,* astrocyte-derived IL15 to microglia) is counterbalanced by intra-cell-type autocrine anti-inflammatory mechanisms (*e.g.*, microglial IL10), suggesting a built-in self-regulation that restrains excessive inflammation. These findings show that glial populations adopt distinct yet complementary functions, balancing propagation of inflammatory signals with mechanisms that prevent overactivation, and the utility of this culture model in uncovering mechanisms that involve multicellular signaling networks.

Despite its advantages, the organotypic slice culture model has limitations. Donor background, genetics, brain region, and tissue handling introduce variability, yet, as shown here, glial responses remain robust and highly reproducible across samples and culture durations. The within-donor paired perturbation framework demonstrated in this study, reduces the inter-individual noise, and also increasing throughput as each donor potentially yields multiple informative conditions. While neuronal survival varies across samples and could affect glial states and variation, we showed that core glial identities remain preserved along culture duration. Moreover, glial responses to stimuli consistently reflected physiologically relevant programs as observed in postmortem brains. Thus, these robust physiological responses along with the within donor paired perturbation design, mitigate the impact of inherent variability and other limitations of this culture system.

Adult cortical resections are routinely obtained during common neurosurgical procedures (brain tumors, epilepsy), and our study joins previous studies^25,28,27^ in demonstrating its feasibility and establishing standardized workflows aligned to standard clinical settings. Cortical tissue is most accessible and advantageous, given its layered organization, consistency of neuronal and glial subtypes across sub-regions, and reduced sampling bias. Here, we integrated samples across sub-regions of the cortex, showing consistent responses. Previous work emphasized neuronal physiology that requires slices with high neuronal survival^26,27^, limiting broader applications. By shifting the focus to glia, shown here to exhibit greater survival and stability within slice cultures, we demonstrate that organotypic slices represent a practical and broadly applicable resource for the scientific community.

Our findings position human organotypic slice cultures as a versatile and translationally relevant platform for functional interrogation of mature glial biology through controlled perturbations within an intact multicellular context. By preserving native brain tissue architecture and mature glial identities across cell types, this model complements existing human iPSC-derived systems while uniquely capturing tissue-level physiological responses driven by multicellular interactions. Looking forward, adapting this platform opens new avenues for studying and modeling glial-driven processes in the human brain, offering broad opportunities to interrogate disease mechanisms and evaluate therapeutic strategies, particularly for age-associated neurodegenerative disorders.

## Extended Data Figures

**Extended Data Figure 1.**
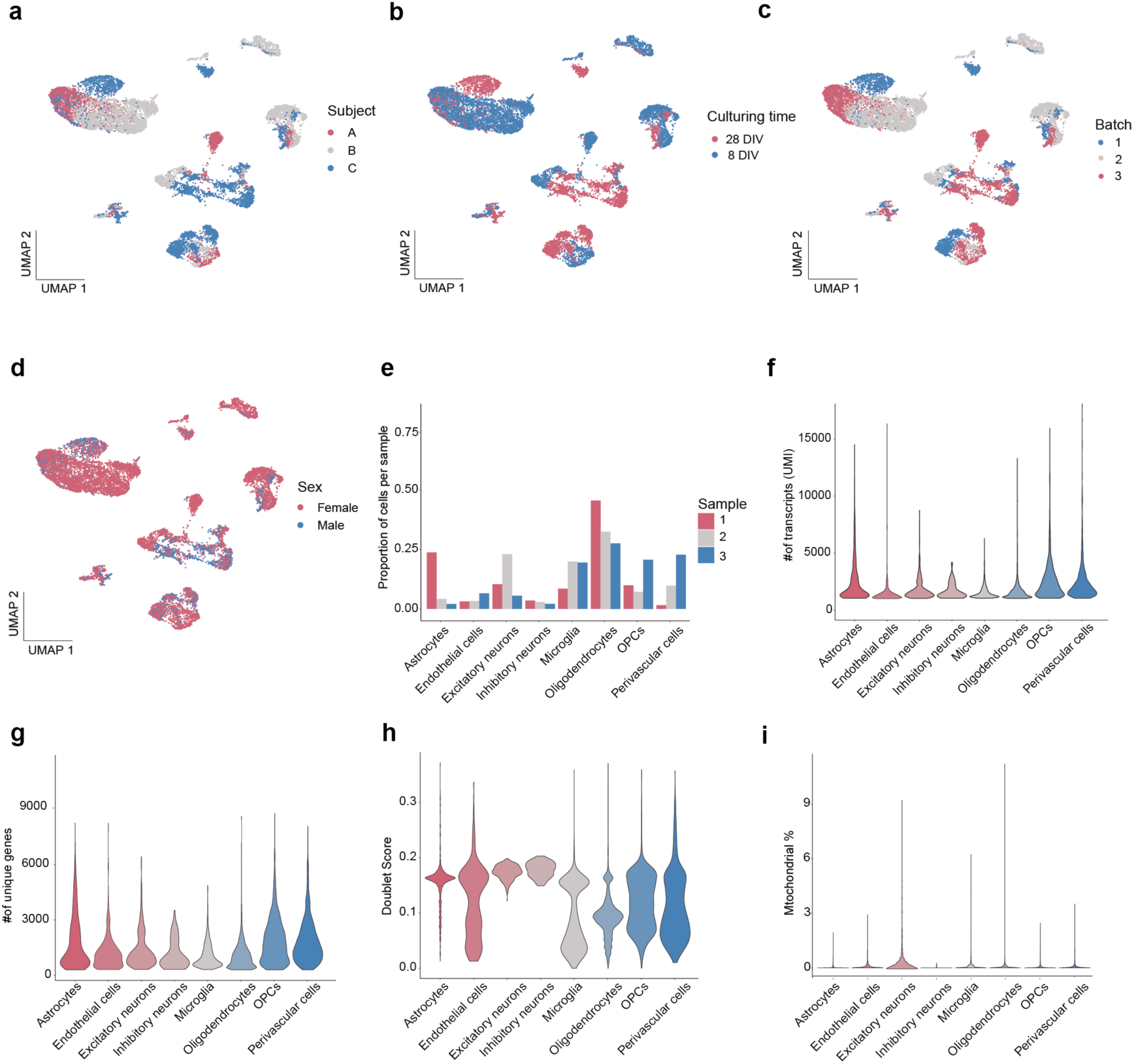
snRNA-seq of slice cultures: Quality controls. **a-d**, UMAP embedding of transcriptional profiles of TNF⍺ treated and control slice cultures. Nuclei (dots) are colored by individuals (**a**), culturing time (**b**), batch (**c**) and sex (**d**). **e,** Cell type proportions per sample across batches. **f-i**, Distribution of transcripts (**f**), number of genes (**g**), doublet score (**h**) and percentage of mitochondrial genes (**i**) per cell type (for n = 11,924 nuclei).

**Extended Data Figure 2.**
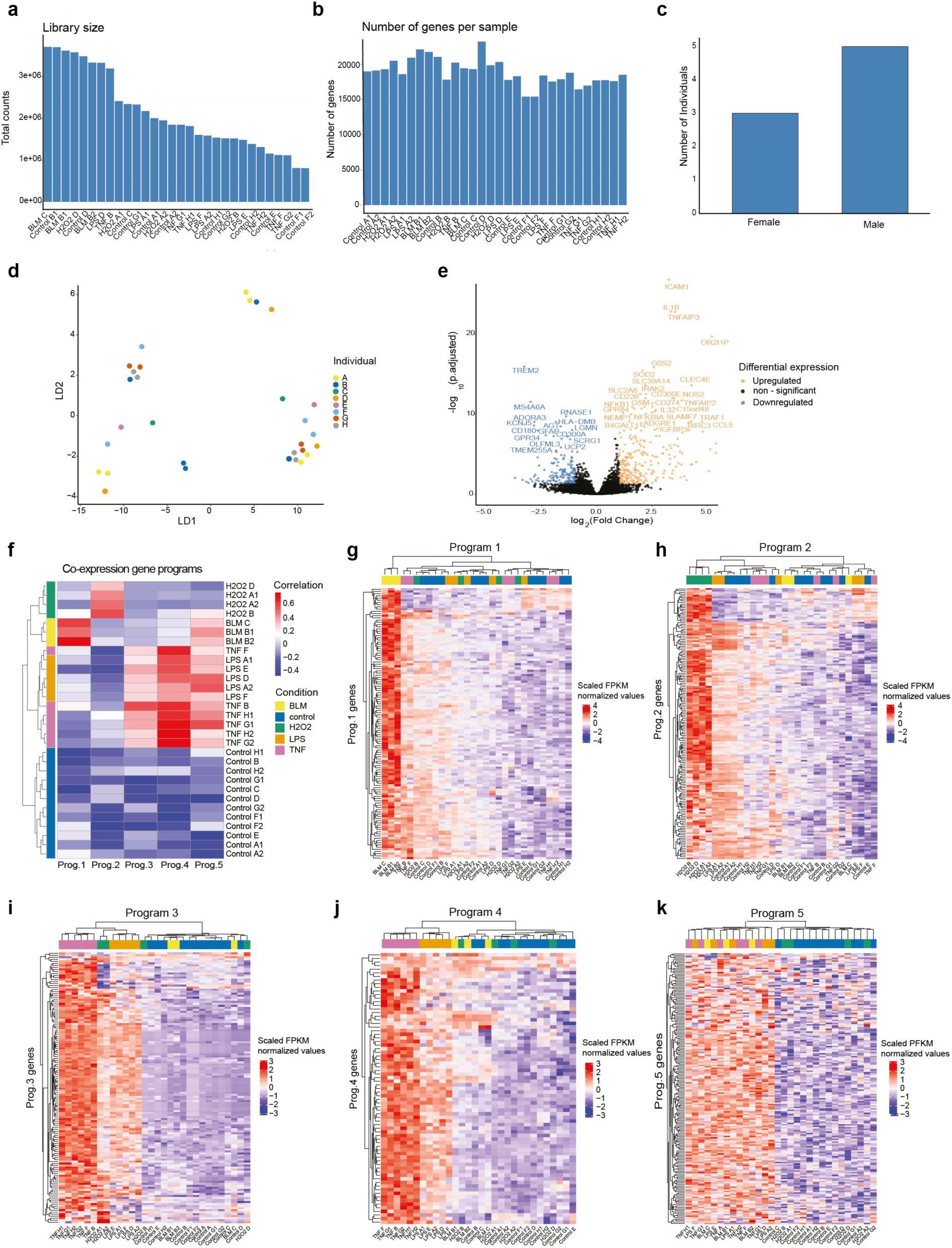
bulk RNAseq of slice cultures: Quality controls. **a**, Distribution of total library size across all samples (n = 30). The y-axis shows sequencing depth per sample. **b**, Number of genes (y-axis) detected above background in each sample (x-axis). **c,** Number of male and female individuals (y-axis) represented per group (n = 8 total individuals). **d,** Embedding of bulk RNA-seq profiles in LDA space, shows differential transcriptional response of slice cultures following TNF⍺, LPS, H₂O₂, BLM (DNA damage)-treatment, colored by individuals. **e**, Differentially expressed genes in response to TNF⍺ in bulk RNAseq data (log_2_ FC > 1.5, adjusted p-value (FDR) < 0.05). **f**, Correlation between WGCNA programs eigengenes and treatments (Pearson correlation, n = 5 programs). **g-k**, Scaled log_2_ normalized expression of genes annotated to the WGCNA programs across samples.

**Extended Data Figure 3.**
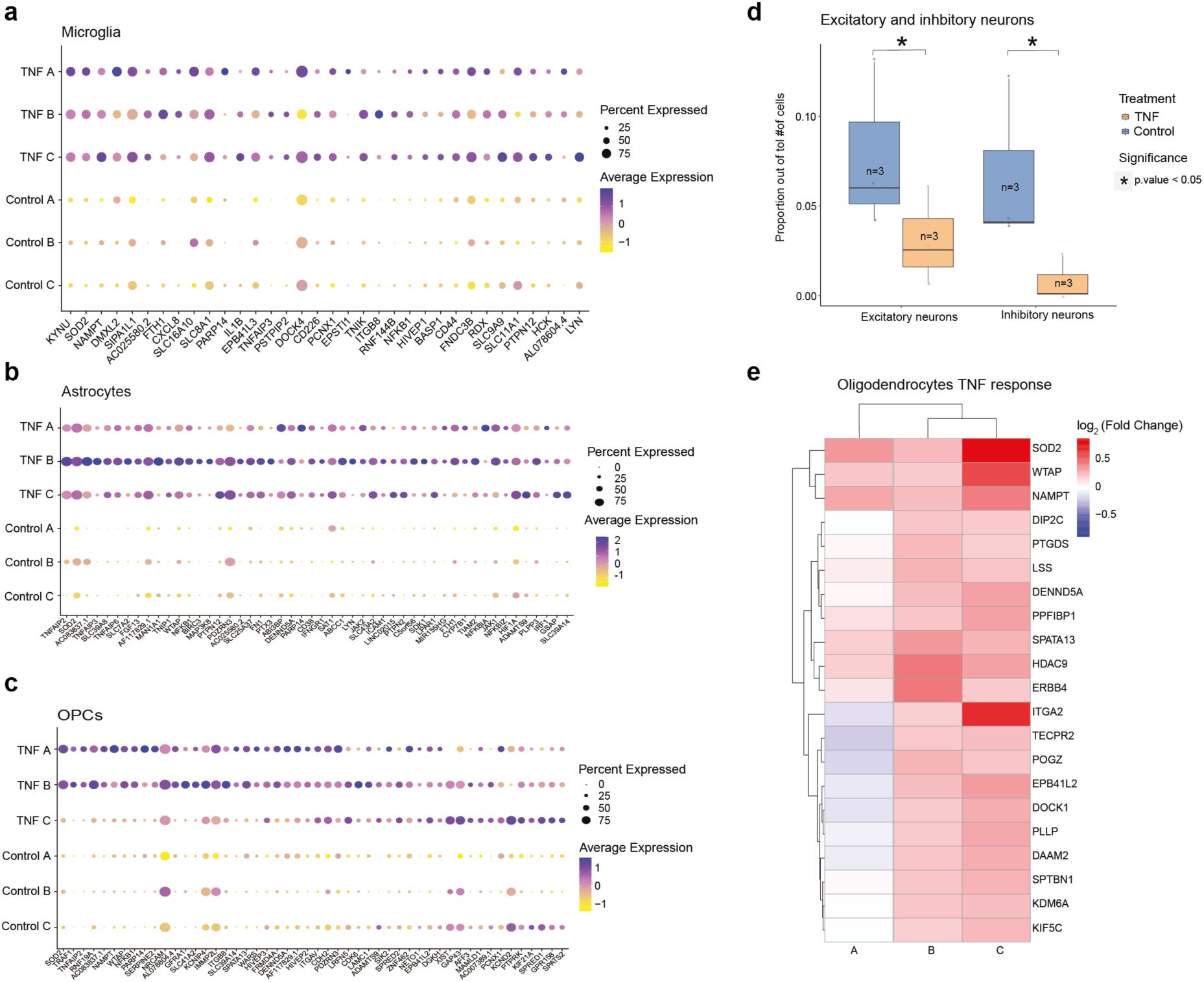
TNF⍺ response across key cell types. **a-c**, Scaled expression of significant TNF⍺-upregulated genes across different samples for microglia (**a**), astrocytes (**b**) and OPCs (**c**). **d**, Proportion of excitatory (left) and inhibitory (right) neurons per sample in TNF-treated and control samples (n=3 per group). P values were calculated using t-tests comparing TNF and control groups. **e**, TNF⍺-induced differential expression in oligodendrocytes. Log_2_ fold-change (TNF⍺ vs. control) of significantly upregulated genes in 8-DIV oligodendrocytes across samples.

**Extended Data Figure 4.**
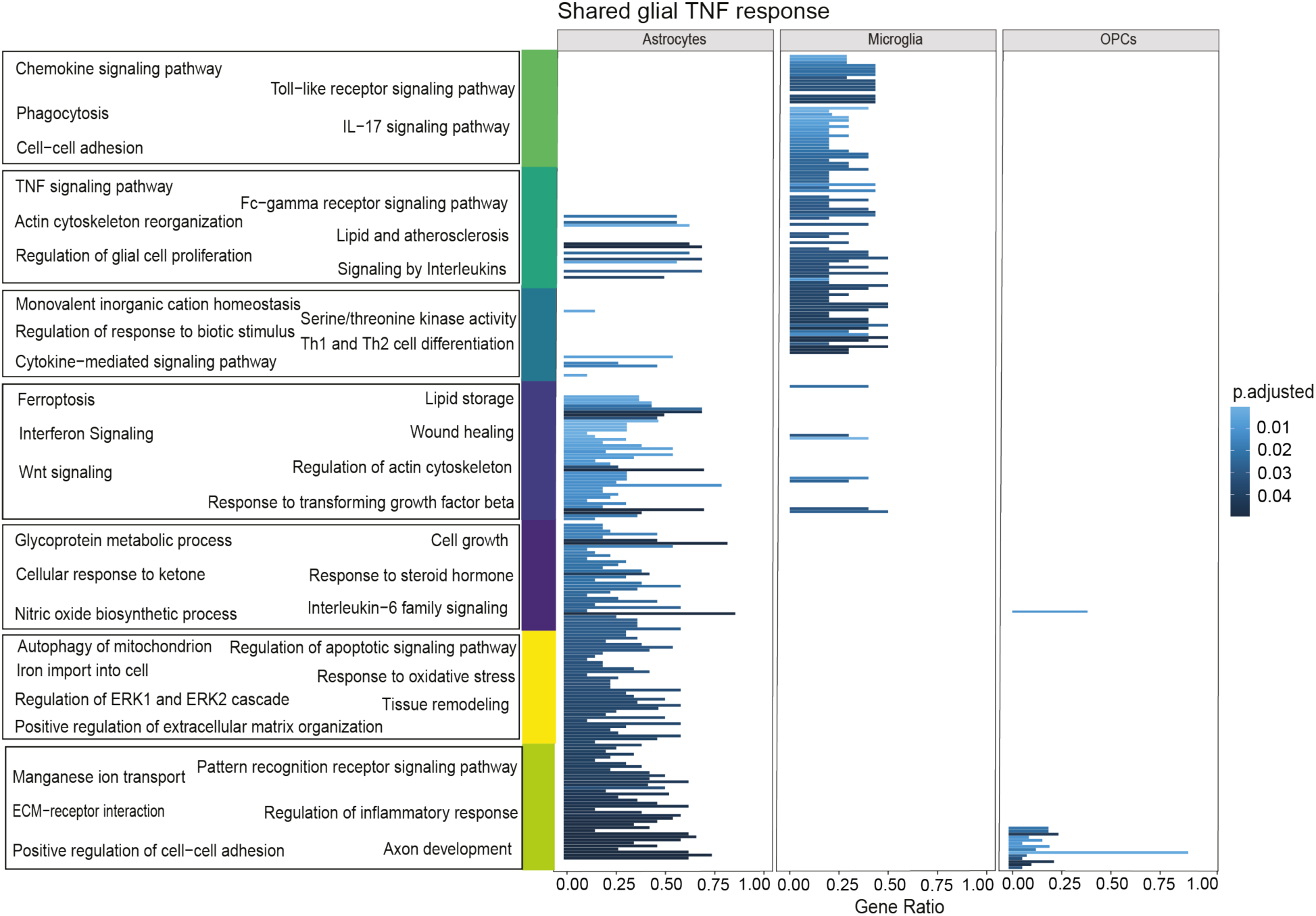
Shared TNF⍺ responses across cell types. Enriched pathways identified across astrocytes, microglia, and OPCs and clustered according to shared genes.

**Extended Data Figure 5.**
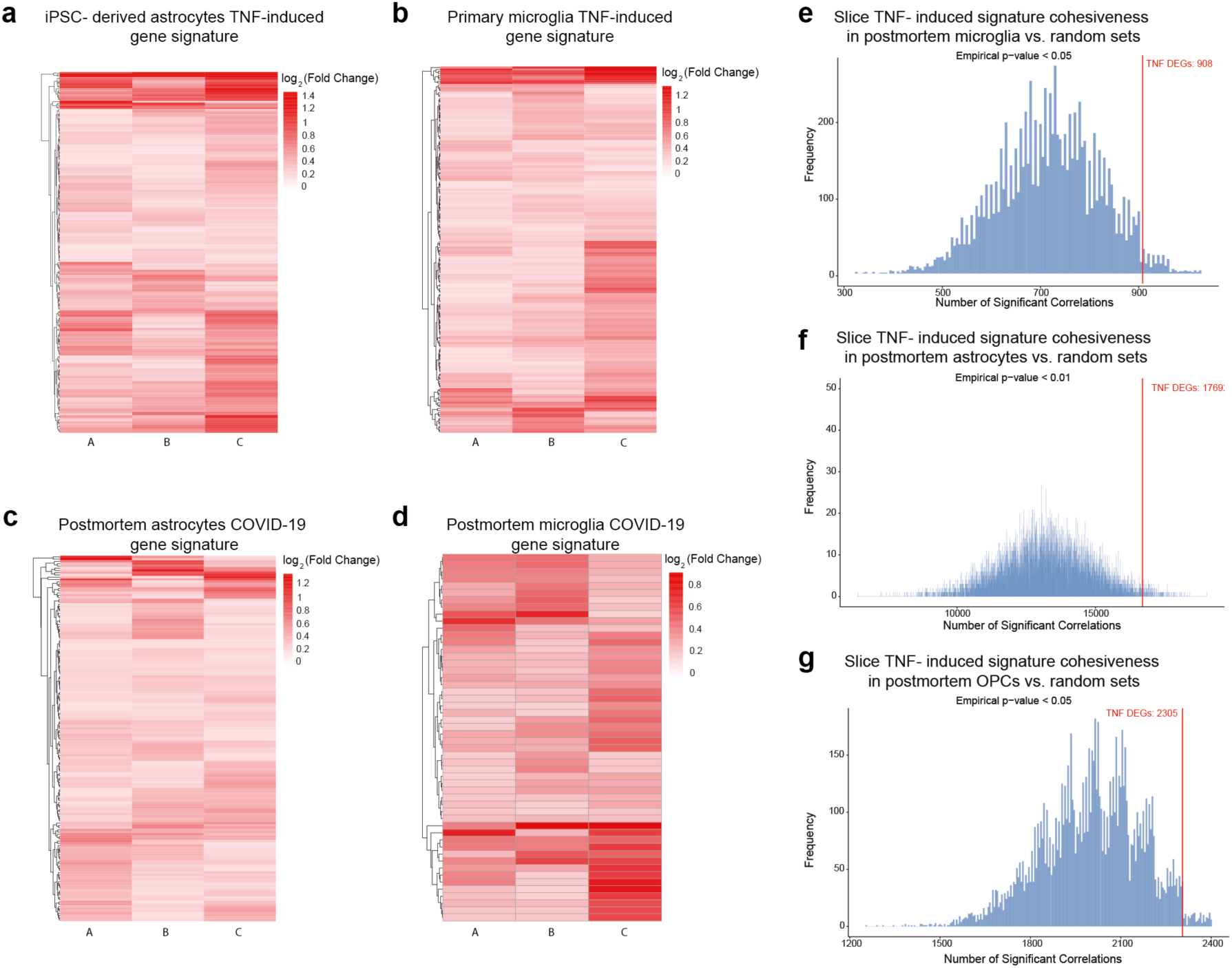
Validation of TNF⍺ response in external datasets. **a**, Differential expression of iPSC astrocytes TNF⍺ signature in slice culture astrocytes. **b.** Differential expression of primary mouse microglia TNF⍺ signature in slice culture microglia. **c-d**, Differential expression of human brain COVID-19 signature in slice culture astrocytes and microglia, respectively. Log_2_ fold-change (TNF⍺ vs. control) of significantly upregulated genes. **e-g,** Slice TNF⍺ induced signature in postmortem glia. Distribution of the number of significant gene-gene correlations in 10,000 randomly selected gene sets matching the size of the TNF⍺ signature for astrocytes (e), microglia (f), and OPCs (g). Slice culture TNF⍺ signature observed co-expression is indicated with red vertical lines. Empirical p-values quantify the probability that random gene sets achieve equal or greater co-expression compared to TNF⍺ signature, computed from Spearman correlations of pseudobulk expression matrices.

## Methods

### Patients and Neurosurgical Procedure

9 patients who underwent resection of deeply located subcortical tumors at the Department of Neurosurgery, Shaare Zedek Medical Center (Jerusalem, Israel) were included in this study (Table 1). All participants provided written informed consent in accordance with protocols approved by the institutional review board (approval numbers: 88-17-SZMC and 159-20-SZMC).

Following dural opening, a corticotomy was performed using low-current bipolar cautery (0.5 mm tip) and an 11-blade scalpel, under guidance of a neuronavigation system, to allow adequate cortical exposure for the resection of the tumor. A Freer elevator was then used to gently disconnect a 1-2 cm² segment of tumor free cortex with adjacent subcortical white matter. All specimens were obtained with meticulous care to minimize mechanical or thermal tissue injury.

### Organotypic brain slice culture procedure

The resected de-identified tissue was transferred into oxygen-enriched (95% oxygen, 5% CO_2_), ice-cold artificial cerebrospinal fluid (aCSF; 110mM choline chloride, 26mM NaHCO_3_, 10mM D-glucose, 11.6mM Na-ascorbate, 7mM MgCl_2_, 3.1mM Na-pyruvate, 2.5mM KCl, 1.25mM NaH_2_PO_4_, and 0.5mM CaCl_2_) and immediately transported to the lab. The tissue was kept in cold, carbogenated aCSF during the tissue block handling and slicing. After pia removal, the tissue block was trimmed to a cubical shape and mounted with glue to the Vibratome stage (Leica, VT1200). 300µm slices were prepared and transferred to room temperature carbogenated aCSF for 30min rest. The slices were then placed on cell culture inserts (Millipore, PICM0RG50) on top of 1.1ml of culturing media – Neurobasal media (Gibco, 21103049) supplemented with 1x serum-free B-27 (Gibco, 17504044), 17mM NaCl, 15 mM Hepes, 13 mM glucose, 2mM GlutaMAX, 1 mM MgSO4 and 1% Pen-Strep) in 6 well plates and incubated at 37°C with 5% CO_2_. Half of the media was changed every 3 days.

Slices were treated for 24 hours with the following compounds: 50nM human TNF⍺ (PeproTech, 300-01A), 100nM LPS (SCBT, sc3535), 20µl/ml Bleomycin sulfate (Fermentek) or for 6 hours with 0.4mM H_2_O_2_. The compounds were added to the culture medium at day 8 or day 21 (DIV), and a small amount of treated culture medium was also added dropwise on top of the slice to form a thin film.

### Immunohistochemistry

Slices were fixed in 4% PFA overnight, followed by permeabilization in 0.5% Triton X-100 for 16hr at 4C. Blocking was performed in 5% BSA for 4 hr in RT. Primary antibodies were incubated for 72 hr at 4°C (1:250 anti-Iba1 Abcam ab5076, 1:1000 anti GFAP Abcam ab7260, 1:5000 anti-Map2, Abcam ab5392) and secondary antibodies incubated overnight. The slices were mounted in ProLong Gold (Invitrogen, P36934) and subjected to imaging with the Leica Stellaris 5 confocal microscope

### Bulk RNA-seq

RNA was extracted with the NucleoSpin RNA kit (MACHEREY-NAGEL) and used for library preparation following the smart-seq2 protocol^79^. Briefly, 10ng of RNA was reverse transcribed (Maxima H minus, Thermo EP0751) using Template Switch Oligo (TSO) to obtain full length cDNA. Tagmentation and library amplification was performed using the Nextera kit (Illumina FC-131-1096). The libraries were quantified using a Qubit and Tapestation and sequenced on Illumina’s Nextseq platform.

### snRNA-seq

The frozen slices were transferred into a dounce homogenizer (Sigma-Aldrich, D8938) with 1.5 mL of lysis buffer. (146 mM Tris HCL pH 7.5, 10 mM NaCl, 3 mM MgCl2, 0.1% NP-40, 40 U/mL of RNAse inhibitors (NEB M0314L). Tissue was gently homogenized while on ice 10 times with pestle A followed by 10 times with pestle B, then transferred to a 15 mL conical tube. A further 3.5 mL of lysis buffer was added to a final volume of 5 mL, left on ice for 5 min, and then centrifuged in a swing bucket rotor at 500 g for 5 min at 4°C. The supernatant was removed, and the pellets were resuspended in 5ml wash buffer (PBS, 1% BSA, 40U/mL of RNAse inhibitor) and centrifuged again at 500 g for 5 mins at 4°C. The supernatant was removed and the pellet was gently resuspended in 250 µL of wash buffer. Nuclei were filtered through a 30 µm MACS Smartstrainer (Miltenyi, 130-098-458) and counted using the LUNA-FL™ Dual Fluorescence Cell Counter (Logos Biosystems) after staining with Acridine Orange/Propidium Iodide Stain (Logos Biosystems, F23001) to differentiate between nuclei and cell debris. The nuclei were partitioned into a nanoliter-scale Gel Bead-In-Emulsion (GEMs) using the 10x Genomics Chromium controller instrument and Single Cell 3’ Reagent Kits v3. Libraries were made following the manufacturer’s protocol.

### Analysis of bulk RNA-seq

Fastq files were mapped to the genome with STAR^80^ and counts per gene were calculated with HTSeq-counts^79^. After preprocessing, raw aligned counts were normalized to fragments per kilobase of transcript per million mapped reads (FPKM, log_10_-transformed) using the countToFPKM (version 1.0) R package, with gene lengths obtained from biomaRt and mean fragment lengths from Picard’s CollectInsertSizeMetrics. Genes with fewer than 10 raw counts across samples were removed prior to analysis. Differential expression between TNF⍺-treated and control samples was performed using DESeq2 (version 1.32.0) with inter-sample differences modeled as covariates. Significance thresholds were defined as FDR-adjusted P. value < 0.05 with log₂ fold change >1.5 (upregulated) or <-1.5 (downregulated). Enrichment of up– and down-regulated genes was performed separately using clusterProfiler (version 3.6.0), testing against KEGG and Gene Ontology (org.Hs.eg.db, version 3.5.0) by hypergeometric test with Benjamini-Hochberg correction (FDR < 0.05). Variance-stabilizing transformation (VST) of counts was applied for downstream analyses. Linear discriminant analysis (LDA) was used to distinguish treatment groups (H₂O₂, BLM, TNF⍺, LPS, and control) while accounting for inter-individual variability, implemented with the MASS (lda function) package in R (version 4.1). Model robustness was evaluated by cross-validation.

For visualization, principal component analysis (PCA) was conducted on VST-transformed counts from TNF⍺-treated samples, restricted to the 500 most variable genes. Batch effects were corrected using the limma package (version 3.50.0, removeBatchEffect function). PCA outputs were visualized with ggplot2 (version 3.4.2).

### WGCNA

Weighted gene co-expression network analysis (WGCNA) was conducted using WGCNA (version 1.70-3) to identify programs of highly correlated genes. Normalized log_10_ (FPKM) expression data for all samples were filtered to retain genes with a variance above the median to reduce noise. A signed adjacency matrix was constructed using a soft-threshold selected based on scale-free topology criteria (R² > 0.85). This matrix was transformed into a topological overlap matrix (TOM), and hierarchical clustering was performed to detect gene programs using dynamic tree cutting (minimum program size = 30), which were subsequently merged (mergeCutHeight = 0.25) based on eigengene dissimilarity. Program eigengenes were correlated with treatment conditions (TNF⍺, LPS, BLM, H₂O₂ and control) using Pearson’s correlation to identify significant and biologically relevant programs (significance threshold |r| > 0.3, adjusted P-value (FDR)< 0.05). Programs highly correlated with individuals were excluded from further analysis.

Hub genes within key programs were identified based on intramodular connectivity (kME > 0.3). Pathway enrichment of significant programs was performed with clusterProfiler (version 4.2.2), using KEGG and Gene Ontology databases (FDR < 0.050, hypergeometric test), as previously described for differential expression results.

### Analysis of snRNA-seq

Fastq files were processed with 10X Cellranger pipeline (v5.0) aligning the reads to the pre-mRNA reference transcriptome.

#### Demultiplexing

Single-cell RNA-seq data were demultiplexed using the Souporcell algorithm^81^ (version 3.6.7), which enables assignment of nuclei to donors in pooled experiments without requiring prior genotype information. Souporcell identifies informative single nucleotide polymorphisms (SNPs) directly from the transcriptome sequencing data, clusters cells by their genotypic profiles, and detects doublets and the presence of ambient RNA. In brief, BAM files from aligned single-cell sequencing reads were used as input for SNP calling, clustering of cell barcodes by inferred genotype, and estimation of doublet rates. Cells were assigned to individual samples based on genotype clusters, and clusters corresponding to doublets or ambiguous assignments were excluded from downstream analysis.

#### Normalization & scaling

Seurat objects were generated for each batch independently. For every nucleus, we quantified the number of genes for which at least one read was mapped. Nuclei with fewer than 300 detected genes and genes detected in fewer than 10 nuclei were excluded. After preprocessing, samples from each batch were merged into a single Seurat object. Gene counts were normalized, variable features were identified, and the data were scaled and centered using the SCTransform Seurat function (version 4.3.0, n.variable.features = 2000).

#### Doublet detection

Doublet scores were assigned to each nucleus using an optimized in-house implementation of DoubletFinder (version 2.0.3), based on the nucleus’ probability of being a doublet, related to the fraction of artificially generated doublet neighbors (parameters: PCs = 1-30, pN = 0.25, pK = 150/(#cells), pANN = False, sct = True). Nuclei with doublet scores above 0.4 were excluded. High-resolution clustering (resolution = 1.5) was applied to identify and remove clusters containing more than 50% doublets.

#### Dimensionality reduction, clustering and visualization

Principal component analysis (using RunPCA function from the Seurat package, npcs = 30) was performed on the scaled expression matrix. Within the top PC space, transcriptionally similar nuclei were clustered together using a graph-based clustering approach. A k-nearest neighbor (k-NN) graph was constructed based on Euclidean distances, refined by the shared overlap of the local neighborhoods using Jaccard similarity (FindNeighbors function, k = 15), and clustered using the Louvain algorithm which iteratively grouped nuclei and located communities in the k-NN graph (using FindClusters function from Seurat, with resolution 0.5). For visualization, nuclei were embedded using Uniform Manifold Approximation and Projection (UMAP), with the top principal components as input (using RunUMAP function from Seurat). Clusters were manually annotated to cell types based on canonical marker gene expression. The distribution of samples within each cluster was examined to eliminate that clusters were driven by batch or other technical effects. Clusters with low-quality cells (low number of genes detected, and missing or low-level cell-type marker genes), as well as doublet clusters expressing markers of multiple cell types, were removed from the analysis.

### Differential expression analysis

Differentially expressed genes (DEGs) between the two conditions within each glial cell type were identified using the MAST test implementation in Seurat’s FindMarkers function (assay set to SCT), accounting for individuals as a latent variable. P-values were adjusted for multiple hypothesis testing using Benjamini-Hochberg’s correction (FDR). Significance was set to FDR-adjusted P-value threshold of 0.05 and a log fold change threshold of 0.25. Volcano plots were generated using ggplot2.

### Pathways enrichment

Pathway enrichment analysis was conducted using clusterProfiler. Gene ontology (GO) enrichment analysis for biological processes (BP), molecular functions (MF), and cellular components (CC), as well as KEGG pathway enrichment for Homo sapiens (’hsa’) and REACTOME pathway enrichment analyses were performed using the enrichGO, enrichKEGG and enrichPathway functions, respectively, with hypergeometric testing and BH FDR < 0.05. Significant pathways were identified based on clustering similar DEGs between pathways using the walktrap algorithm and kappa score implemented in the igraph package (version 1.2.11). Enriched pathways were visualized with ggplot2.

### Cross cell-type comparisons

To identify shared transcriptional responses across glial cell types, differentially upregulated genes (DEGs) were selected based on adjusted p-values (FDR < 0.05) and fold change thresholds (log₂FC>0.25). Genes that were significantly upregulated in more than a single glial cell type in response to TNF⍺ treatment were clustered based on expression similarity. Hierarchical clustering was applied to group genes based on shared expression patterns across glial cell types, prioritizing genes consistently upregulated in all selected cell types, followed by those shared across specific groups of cell types. Based on the clustering results, top genes shared between all major cell types were selected and visualized using pheatmap (version 1.0.12).

Shared pathways were defined by performing pathway enrichment analysis for each cell type’s TNF⍺ signature using EnrichGO and EnrichKEGG tools, testing for overrepresentation in curated gene sets. Enriched pathways from all cell types were then clustered by shared gene content using network-based clustering. A bipartite graph was constructed where nodes represent pathways connected if they shared DEGs, and pathway clusters were identified using Louvain community detection and kappa statistics. Resulting pathway clusters were compared and visualized across the three cell types with ggplot2.

### Validations

TNF⍺-induced signatures were validated in previously published bulk RNA-seq datasets of mouse microglia^67^ and iPSC-derived astrocytes^68^. Raw count data was processed using the DESeq2 package, following the analytical parameters and workflows described in the respective publications. Differentially upregulated genes (log₂FC > 1.5, FDR < 0.05) were identified and filtered to include those expressed in more than 10% of the microglia or astrocyte populations, respectively. Validation was further extended to COVID-19 associated signatures by retrieving astrocyte– and microglia-specific DEGs from postmortem COVID-19 brain transcriptomic datasets^69^. Genes overlapping with those expressed in at least 10% of cells in the organotypic slice culture were retained. A signature score was calculated using the AddModuleScore Seurat function to quantify the TNF⍺-induced as well as COVID-19 transcriptional signature in the slice culture experimental data.

### Statistical analysis

Treatment effects on signature scores were assessed using linear mixed-effects models including treatment as a fixed effect and individuals as a random intercept to account for inter-sample variability. Statistical significance of treatment terms was determined using likelihood ratio tests comparing full models with reduced models lacking the treatment effect. P-values were adjusted by Benjamini-Hochberg. Analyses were conducted using the lme4 (version 1.1-32) and lmtest (version 0.9-39) R packages.

### Postmortem pseudobulk analysis

Pseudobulk RNA-sequencing profiles were generated from 437 postmortem human brains^10^ and from TNF⍺-treated human organotypic brain cultures. For the postmortem dataset, cell type-specific matrices were obtained by averaging log-normalized expression values across all nuclei annotated to each major cell type per individual, ensuring that expression values reflected population-level transcriptional activity. For cross-dataset comparisons, expression was restricted to genes present in both sources prior to analysis. To assess transcriptional similarity, postmortem profiles from 287 individuals classified as healthy aging based on their probability of assignment to the healthy trajectory in pseudotime analysis (see Green et al.^10^) were averaged by cell type and correlated with each corresponding organotypic brain sample using Pearson correlation. Differentially expressed genes (DEGs) induced by TNF⍺ were identified in organotypic astrocytes, microglia, and oligodendrocyte progenitor cells (OPCs) using Seurat’s FindMarkers function (log₂ fold change > 0.25, adjusted p. value (FDR) < 0.05; see Differential expression analysis section). Gene lists were further intersected with the postmortem dataset and functionally annotated.

To quantify TNF⍺ signature cohesion within each cell type, pairwise gene-gene Spearman correlations were computed from pseudobulk expression of the slice culture TNF⍺ signature, and the upper triangle of the resulting correlation matrices was extracted to obtain unique gene-gene pairs. Observed correlation distributions were compared against 10,000 random-size gene sets of equal size, producing empirical p-values representing the probability of obtaining an equal or higher level of coexpression by chance.

To assess inter-cell type coordination, log-transformed mean signature scores were calculated for the slice culture TNF⍺ signature in each cell type per individual and correlated pairwise between astrocytes, microglia, and OPCs. Scatter plots of these scores were colored by the diagnostic group (healthy aging, Alzheimer’s disease, or non-AD aging), with regression lines and associated correlation coefficients and p-values overlaid.

For astrocytes, mean expression scores were also computed for each predefined functional gene set and correlated within each other and the TNF⍺ signature score using Spearman correlation, allowing identification of pathways whose activity tracked with TNF⍺-induced changes. Correlation coefficients were visualized as heatmaps and scatter plots to highlight patterns of co-regulation.

All statistical analyses were performed using stats package (version 4.1.2) for correlation inference and resampling procedures for empirical significance testing; multiple p-values were FDR-adjusted by the Benjamini-Hochberg method. Data visualization was carried out with ggplot2 and ComplexHeatmap (version 2.0.10).

### Validations in iPSC-derived astrocytes

iPSC cell lines used in this study were obtained in collaboration with the Collaborative Center for XDP (CCXDP), at MGH (Boston, MA, USA) and were generated from neurologically-normal human fibroblasts as previously described1,2.

iPSC-derived astrocytes were differentiated following a modified protocol^82^. On day 0, iPSCs at 80-90% confluency were dissociated with Accutase (Life Technologies, 00-4555-56), and 6 × 10^5^ cells/cm^2^ were plated in Essential 6 medium (Thermo Fisher Scientific, A1516401) with 10 µM ROCK inhibitor (StemCell Technologies, Y-27632). On days 1-2, cells were fed with Essential 6 medium supplemented with 2 µM XAV 939 (Tocris Bioscience, 3748), 100 nM LDN (Tocris Bioscience, 6053), and 10 µM SB (Tocris Bioscience, 1614). On days 3-9, cells were fed with Essential 6 medium supplemented with 100nM LDN and 10 µM SB. On days 10-16, cells were fed with medium containing 50% Neurobasal medium (Thermo Fisher Scientific, 21103049), 50% DMEM/F12 medium (Thermo Fisher Scientific, 11320033), 2% B27 supplement (Thermo Fisher Scientific, 17504044), 1% N2 supplement (Thermo Fisher Scientific, 17502048), and 1% penicillin-streptomycin (Thermo Fisher Scientific, 15070063) (N2/B27 medium). On day 16, cells were dissociated with Accutase, and 2 × 10^5^ cells/cm^2^ were plated on Matrigel-coated 6-well plates in N2/B27 medium containing 10 µM ROCK inhibitor. On day 17, cells were fed with N2/B27 medium. On day 18, cells were transduced with 2 µg FUW-NFIA and 2 µg FUW M2-rtTA lentiviruses in N2/B27 medium. Lentivirus packaging was performed by the Boston Children’s Hospital Viral Core Facility. On days 19-24, cells were fed with N2/B27 medium containing 1 µg/mL doxycycline hyclate (Stemcell Technologies, 72742). On days 25-45, cells were propagated with media changes of astrocyte medium (ScienCell, 1801) with 2% FBS every two days and passaged at 80-90% confluency (complete AM). On days 45-60, cells were switched to astrocyte media with 0% FBS and grown as on days 25-45.

iPSC-derived astrocytes were seeded at 30,000 cells per well onto 24 well plates, allowed to rest for 3 days, and then stimulated for 24 hours with 50 ng/mL human TNF (Gibco, PHC3016), 100 ng/mL human IL-1β (Gibco, PHC0815), or both in FBS-free astrocyte medium. After 24 hours, media was removed, cells were lysed in RLT buffer, and RNA was isolated using the QIAGEN RNeasy Mini kit (QIAGEN, 74106). Libraries were prepared following SMARTSeq2 protocol^83^ and sequenced at the Broad Institute Technology Labs and the Broad Genomics Platform.

### Comparison of iPSC-derived astrocytes and slice astrocytes

Bulk RNA-seq data from iPSC samples were normalized using DESeq2, with batch correction performed via limma’s removeBatchEffect. Organotypic astrocytes pseudobulk profiles were generated by aggregating single cell counts from astrocyte subpopulations. DEG-derived gene signatures from TNF⍺-treated organotypic astrocytes were used to calculate signature scores in bulk samples by averaging expression levels following batch correction. Statistical significance of signature score differences across conditions was assessed by pairwise t-tests with Benjamini-Hochberg correction. PCA was performed across datasets, projecting organotypic astrocyte pseudobulk signatures onto iPSC PC embedding using weighted sums of top 500 positive and negative PC genes. Results were visualized in PCA plots combining both datasets using ggplot2.

### Cell-cell signaling network analysis

Cell-cell communication networks were assessed with the MultiNicheNet package (version 1.0.3) with default parameters to compare ligand-receptor interactions between TNF⍺-treated and control samples across the three main glial cell subtypes. Differential interaction analysis was performed using pseudo-bulk aggregation followed by edgeR-based modeling to identify ligand-receptor pairs with significant expression changes (FDR < 0.05, |log₂FC| > 0.25), while accounting for inter-sample variability. Prioritized interactions were further refined using ligand activity scores derived from NicheNet-v2 regulatory networks and expression prevalence across conditions. Key interactions were visualized via circos plots to map top 50 global communication patterns between glial subtypes comparing TNF⍺ interactions to control. Selected ligand-receptor-target interactions and LR interactions were visualized with ggplot2.

## Data availability

The sequencing datasets generated in this study will be available in the database of Genotypes and Phenotypes (dbGaP)

## Code availability

The code base used in to derive the main figures and analysis conducted as part of this study is available at GitHub (https://github.com/naomihabiblab/HumanOrganotypicTNF).

## Acknowledgements

We thank the individuals who donated brain tissue samples to research through. This work was supported by the Israel Science Foundation (ISF) research grant no. 1709/19, the European Research Council grant 853409 (N.H.); The Chan Zuckerberg Initiative (CZI) Collaborative Pairs grant CP2-1-0000000058 and the Myers Foundation (N.H. and F.J.Q.);

## Author contributions

N.H., M.A., I.S. conceived and designed the study, data interpretation and wrote the manuscript with critical comments from all the co-authors; T.S., I.P. performed and facilitated the collection of specimens from brain surgeries, with help from H.S; M.A. established protocols and with I.S. performed all experiments involving organotypic slice cultures, and generation of the transcriptomic and immunohistochemistry data; G.P., Z.L., and F.J.Q. performed the experiments in iPSC-derived astrocytes. I.S. performed the computational analyses with help from M.A., and guidance from N.H.

## Supplementary Tables and Movie

**Supplementary Table 1. Samples description:** Characteristics of all human tissue samples used in this study (tumor type, brain region, sex and age).

**Supplementary Table 2. Bulk transcriptomic signatures across stimuli:** WGCNA analysis per stimuli and associated pathways (for TNF, LPS, BLM and H2O2).

**Supplementary Table 3. Bulk transcriptomic response to TNF:** Differentially expressed genes and enriched pathways following TNF stimulus.

**Supplementary Table 4. Cell-type specific transcriptomic response to TNF:** snRNA-seq differential signatures and enriched pathways in: microglia, astrocytes, OPCs.

**Supplementary Table 5. MultiNicheNet signaling network analysis:** Prioritization table for top 70 interactions under control and TNF.

**Supplementary Movie. MAP2 staining of neurons in 3D.** Immunohistochemistry with anti-MAP2 antibody showing neurons across the depth of a organotypic adult human brain slice culture at 8DIV, 20x magnification.

