## Supplementary Table 1 for "Adult human *ex vivo* brain slices for dissecting glial biology and multicellular communication"

### List of samples included in the study

| Age | Sex | Area | Type of tumor | Bulk RNA-seq | snuc-RNA-seq |
|---|---|---|---|---|---|
| 22 | M | Frontal | Glioma | x | |
| 67 | M | Frontal | Metastatic Carcinoma | x | |
| 63 | F | Temporal | Glioma | x | |
| 45 | F | Frontal | Glioma | x | x |
| 43 | M | Frontal | Glioma | x | x |
| 68 | F | Parietal | Glioma | x | x |
| 64 | M | Frontal | Brain Metastasis | x | |
| 69 | M | Frontal | Meningioma | x | |
| 64 | F | Frontal | Metastatic Carcinoma | | x |
